# blockCV: an R package for generating spatially or environmentally separated folds for k-fold cross-validation of species distribution models

**DOI:** 10.1101/357798

**Authors:** Roozbeh Valavi, Jane Elith, José J. Lahoz-Monfort, Gurutzeta Guillera-Arroita

**Affiliations:** School of Biosciences, University of Melbourne, Parkville, Vic. 3010, Australia

**Keywords:** block cross-validation, environmental blocking, model evaluation, structured environment, spatial autocorrelation, spatial blocking, spatial leave-one-out, species distribution modelling

## Abstract

1. When applied to structured data, conventional random cross-validation techniques can lead to underestimation of prediction error, and may result in inappropriate model selection.
2. We present the R package **blockCV**, a new toolbox for cross-validation of species distribution modelling.
3. The package can generate spatially or environmentally separated folds. It includes tools to measure spatial autocorrelation ranges in candidate covariates, providing the user with insights into the spatial structure in these data. It also offers interactive graphical capabilities for creating spatial blocks and exploring data folds.
4. Package **blockCV** enables modellers to more easily implement a range of evaluation approaches. It will help the modelling community learn more about the impacts of evaluation approaches on our understanding of predictive performance of species distribution models.

## Introduction

Species distribution modelling (SDM) is a popular tool in Ecology, in part because it is able to produce spatial predictions of species distributions. An important component of the modelling process is evaluation of the output. Is it fit for purpose? Are some models more suited to the nominated end-use than others? Here we introduce a software package relevant to these questions.

When evaluating SDM performance, it is common to test how well predictions match observations at a set of locations. Whilst early applications of SDM tended to focus on statistical measures of model fit on the data used to fit the model, attention has gradually shifted towards testing on independent data. Since fully independent data (such as new surveys) are rarely available, a common approach involves sub-sampling the data available for modelling. In ecology, this usually involves splitting the data into subsets for training and testing (also known as calibration and validation, respectively). Training data are used for fitting the model, and testing data, for evaluating performance of the trained model. This is termed cross-validation, with variants including simple random splits, repeated random splits, or k-fold cross-validation (Box 1).

In recent years, some discussion has focussed on the best way to allocate data to train and test datasets. Should allocation be random, or should structure be introduced? This cannot be answered independent of knowledge of the data and the modelling task. However, there are many situations that motivate non-random allocation of sites to train and test sets. SDMs are now commonly used to predict to new times or places, so tests of their ability to predict to temporally, environmentally or geographically separated subsets of data may be the most useful indicators of their predictive performance for these tasks (Bahn & McGill, 2012; Wenger & Olden, 2012; Roberts et al., 2017). Ecological data are often autocorrelated – i.e. observations close to each other (in space or time) are more similar than distant ones (Legendre, 1993). In species distribution modelling this is true of the response (the species data) and the predictor variables, and might result from biotic or abiotic processes. Spatially-separated training and testing datasets can help test whether the model performs as well in nearby locations as it does in more distant places – if it does not, structure might not be properly accounted for in the model, the model might be over-fitted to the training data, or important predictor variables might be missing altogether (Dormann et al., 2007; Roberts et al., 2017). The package we introduce here, written in the R programming language (R Development Core Team, 2017) provides useful tools for achieving this.

### The blockCV R package: overview

Other software packages do exist for creating segregated datasets for cross-validation − for instance, R packages **sperrorest** (Brenning, 2012) and **ENMeval** (Muscarella et al., 2014) and the Python-based ArcGIS **SDMtoolbox** (Brown et al., 2017). These have various relevant features, but as users of SDMs we found them limited in their applicability to distribution modelling across typical nuances of species data and analytical aims. Package **blockCV** aims to fill that gap.

In a nutshell, package **blockCV** provides functions to build train and test data sets using three general strategies, described in detail below: *buffers*, *spatial* and *environmental* blocks. It offers several options to construct those blocks and to allocate them to cross-validation folds (see definitions, Box 1). It includes a function that applies geostatistical techniques to investigate the existing level of spatial autocorrelation in the chosen predictor variables, to inform the choice of block and buffer size. In addition, visualization tools further aid selection of block size and provide understanding of the spread of species data across generated folds. The package has been written with *species distribution modelling* in mind, though we have kept the output general enough that it is likely to be useful more generally. The functions allow for a number of common usage scenarios, including presence-absence and presence background species data (Box 1), rare and common species, and raster data for predictor variables.

The generated output is stored in lists that identify allocation of locations to train or test data and, where relevant, to cross-validation folds. These can be directly input to any species modelling workflow in R, and formats for the widely-used species distribution modelling package biomod2 (Thuiller et al., 2017) are also included. With the package, we provide a vignette and example data to demonstrate use of these functions in modelling. The following sections describe the functionalities of the package in more detail.

#### Box 1: Cross-validation, blocks, folds and species data − definitions

##### Cross-validation and folds

Cross-validation (cv) is a technique for evaluating predictive models. It partitions the data into k parts (*folds*), using one part for testing and the remaining (k-1 folds) for model fitting. In k-fold cv the process is iterated until all the folds have been used for testing. If k is equal to the number of records, it is called n-fold or leave-one-out (LOO) cross-validation (Hastie et al., 2009, p. 241).

##### Blocks and folds

*Blocks* are units of geographical area (e.g. rectangles, spatial polygons and buffers of specific distance) or environmental clusters. In these units, all species data are treated together – for instance, allocated to the same fold of cv. Several blocks could be allocated to one cv fold.

##### Species data

Presence-only data are locations where a species was observed. These can be coupled with a large sample of points across the landscape, known as background data (Renner et al., 2015). Presence-absence data include both the locations where the species has been observed and where it has been searched for, but not found (Guillera-Arroita et al., 2015). Following common convention, our code expects presence data to be represented by a 1 and absence or background by a 0.

## Initial considerations

In blocking methods, decisions need to be made about how to group the blocks into folds for cross-validation (Box 1). The practical implementation of this “blocks into folds” step is affected by the species data, as described in the following sections. The package currently explicitly deals with the two main types of species response data (Box 1): presence-only (with background points, where relevant) and presence-absence data, though the methods implemented for presence-only data could be applied to any type of data where each site is the unit of interest.

The package assumes that species spatial data and raster predictor variables have the same projection and similar extent –i.e. the rasters at least extend across all species data. By default, the package creates blocks according to the extent and shape of the study area, based on the rasters. Alternatively, the spatial blocks can be masked based on species spatial data. This is especially useful when the species data are not evenly dispersed across the whole region e.g. for rare species.

Spatial data sometimes have coordinate reference systems in degrees. Whilst the most satisfactory approach is for the user to first convert these to metric reference systems (e.g. UTMs), the package provides alternatives that handle data in degrees. For instance, those blocking strategies relying on distance (size of the blocks or buffers) use an (optional) scaling factor to convert degree to metres. Computations (described later) regarding spatial autocorrelation in raster data with geographic reference system use the great circle distance (the shortest distance between two points on the surface of a sphere; Longley et al., 2015, p. 305).

## Blocking strategies

**BlockCV** supports three strategies to separate train and test data: spatial blocking, spatial buffering, and environmental blocking, each explained in detail below. We first explain the strategies, then discuss approaches for choosing block or radius size.

### Spatial blocking

A general strategy to account for spatial autocorrelation when evaluating models is to split data spatially into blocks. Package **blockCV** provides several methods to create spatial blocks (Fig. 1). One is to build square spatial blocks of a specified size (i.e. width). The spatial position of these blocks can be shifted vertically or horizontally, allowing assessment of the sensitivity of model evaluation metrics to specific block arrangements. The package also allows division of the study area into vertical or horizontal bins of a given height or width (rectangle blocks, Fig. 1), as used by Wenger & Olden (2012) and Bahn & McGill (2012) respectively. Finally, the blocks can be specified by a user-defined spatial polygon layer (Fig. 1).

**Fig 1.**
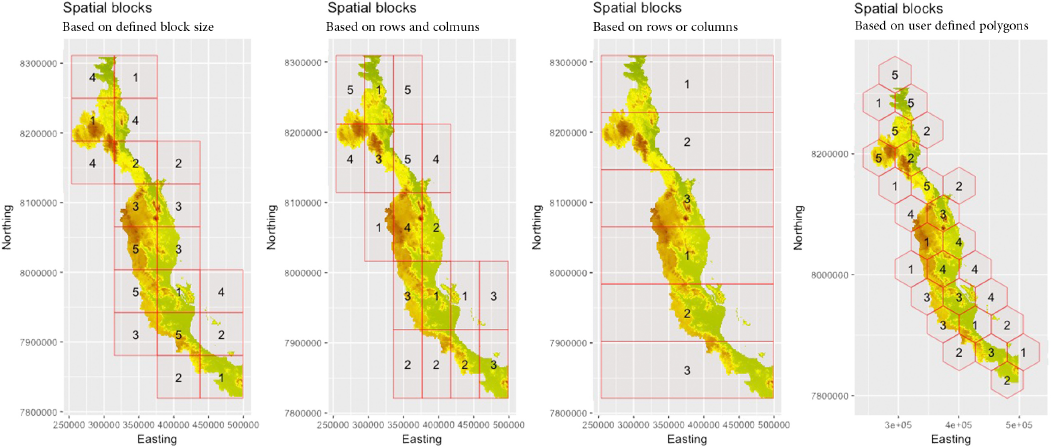
Illustration of different spatial blocking strategies. Blocks are outlined in red. The numbers in the blocksare fold numbers, showing allocation of blocks to folds.

The package allows allocation of blocks to folds (Box 1) both randomly and systematically, and this is one of the key important steps for species modelling because species data are rarely evenly dispersed over landscapes. When random selection of folds is chosen,constraints can be set to avoid folds with little or no presence or (where relevant) absence data. It is also possible to find block-to-fold allocations that achieve most even spread of species data across folds (e.g., a similar number of presence and absence records in each fold). In systematic allocation, blocks are numbered and assigned to folds sequentially. As explained later, interactive tools are provided for tabulating and visualising the placement of folds and distribution of species data over folds.

### Buffering

The buffering strategy generates spatially separated train and test folds by considering buffers of specified radius around each observation point (Le Rest et al., 2014). This approach is related to leave-one-out cross-validation (Box 1), and can be desirable if a user wants to ensure that no test data abuts training data. In this case there is no need to distinguish between blocks (buffered points) and folds, because each left-out point is equivalent to a fold. In the following description, we envisage the method from the viewpoint of the test data. The approach varies slightly with the type of species data available (specified by *spDataType* argument).

For *presence-absence* data, folds are created based on all records, both presences and absences. Each target observation (presence or absence) forms a test point; all presence and absence points other than the target point within the buffer are ignored. The training set comprises all presences and absences outside the buffer.

When working with *presence-background* data, test folds are determined by the presence data. Given a target presence point to be predicted, a buffer is defined around it using the specified radius (range). By default, the testing fold comprises the target presence point and all background points within the buffer (Fig. 2). Since some modellers may wish to deal only with presence points, there is an option (addBG = FALSE) for ignoring background data. Any non-target presence points inside the buffer are excluded from both the training and testing sets. All points (presence and background) outside the buffer are used as the training set. The method cycles through all the presence data, each time allocating one point for testing, so the number of folds is equal to the number of presence points in the dataset.

**Fig. 2.**
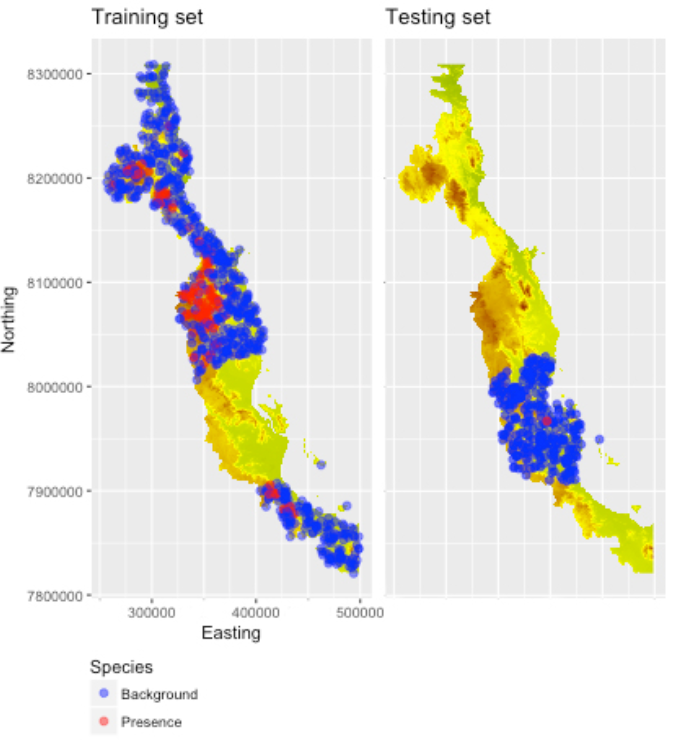
The buffering fold with the target point (presence) and background points in the testing fold

A “species” argument optionally points to the column with species data. By default, it is set to null – this is the relevant setting for presence-only data (with no absence or background points) or other types of response data (e.g. multi-class points for classifying remotely sensed imagery). For presence-absence or presence-background data it should be used to direct the code to the column of data containing the species data (0s and 1s, see Box 1).

~~~
*# example code of buffering with presence-background data*
bf2 <- **buffering**(speciesData= pb_data, *# presence-background data*
                                    species= "Species", *# columns with 0s and 1s*
                                    theRange= 68000, *# radius (m) the of buffer*
                                    spDataType = "PB", *# presence-background data type*
                                    addBG = TRUE) *# add background data to testing folds*
~~~

### Environmental blocking

This strategy uses clustering methods (k-means; Hartigan & Wong, 1979) to specify sets of similar environmental conditions based on the input covariates and the chosen number of clusters in (possibly multivariate) environmental space. Species data within each cluster are assigned to a fold, so the number of folds is by default equal to number of clusters chosen by the user. This algorithm only makes sense with continuous raster data (i.e. categorical covariates like vegetation classes should be excluded).

The clustering can be based on all raster cells or only on values at the species points. When based on the rasters, the clusters will be consistent throughout the region and across all species being considered in that region. However, this does not guarantee that all clusters contain species records, especially when species data are not dispersed across all environments. So, the resulting folds in practice might be fewer than the specified k. Alternatively, the clustering can be done based only on the values of the predictors at the species presence and absence points. In this case, the number of the folds will always be the same as k.

### Choosing block size

One of the challenges of using spatial blocks or buffered cross-validation is choosing the optimal size of blocks or buffers (Trachsel & Telford, 2016). The spatial autocorrelation range in model residuals has been used to define the optimal separation distance between train and test sets (Telford & Birks, 2009; Trachsel & Telford, 2016; Roberts et al., 2017). This is the range over which residuals are independent, and can be characterized using the empirical variogram, a fundamental geostatistical tool for measuring spatial autocorrelation. The empirical variogram describes the structure of spatial autocorrelation by measuring variability between all pairs of points (O’Sullivan & Unwin, 2010).

Having the residuals, however, implies having fitted the model already. To apply block CV methods, we need to make considerations about potential autocorrelation *prior* to model fitting. One option suggested for presence-absence data is to fit a variogram to the raw species data (Roberts et al., 2017 and see Bio et al., 2002) and use the resulting distance from the analysis as block size. Alternatively, to support a first choice of block size, prior to any model fitting, package **blockCV** allows the user to look at the existing autocorrelation in the predictors, as an indication of landscape spatial structure (Fig. 3). The function works by automatically fitting variogram models to each continuous raster and finding the effective range of spatial autocorrelation. A number of random points (5000 by default) is taken from each input raster and parallel processing is used to speed up the computation. For the sake of simplicity, we used isotropic variogram (non-directional) and assumed that the data met the necessary geostatistical criteria e.g. stationarity (having constant variance). The variogram fitting procedure uses the automap package (Hiemstra et al., 2009). Output plots show the spatial autocorrelation ranges of input raster covariates and the spatial block that has been created based on median of these ranges (Fig. 3).

**Fig. 3.**
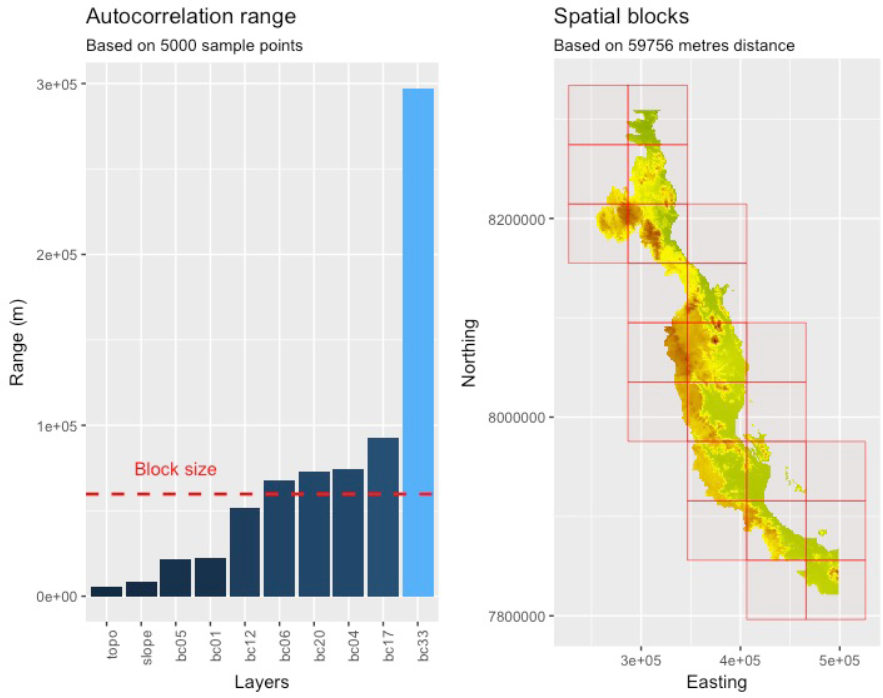
Output graph from the package **blockCV** showing: (a) spatial autocorrelation ranges in input covariates,and (b) corresponding spatial blocks (the selected block size is based on median SAC range across all input data)

### Interactive visualisation tools

Package **blockCV** provides two visualisation tools, developed as local web applications using R-package shiny (Chang et al., 2017). Through a user interface, these tools (1) allow for graphical exploration of the generated folds (the *foldExplorer* tool) and (2) assist the selection of a suitable spatial block size (the *rangeExplorer* tool). The *foldExplorer* tool displays a map where folds are overlaid, and provides a summary of the number of records in each fold. This helps the user assess the adequacy of the distribution of train and test folds throughout the study area. The tool is available for all three blocking strategies. The *rangeExplorer* tool (Fig. 4) allows the user to interactively change the size of spatial blocks, visualise the blocks and assess the impact of block size on the number and arrangement of blocks in the landscape (and optionally on the distribution of species data in those blocks).

**Fig. 4.**
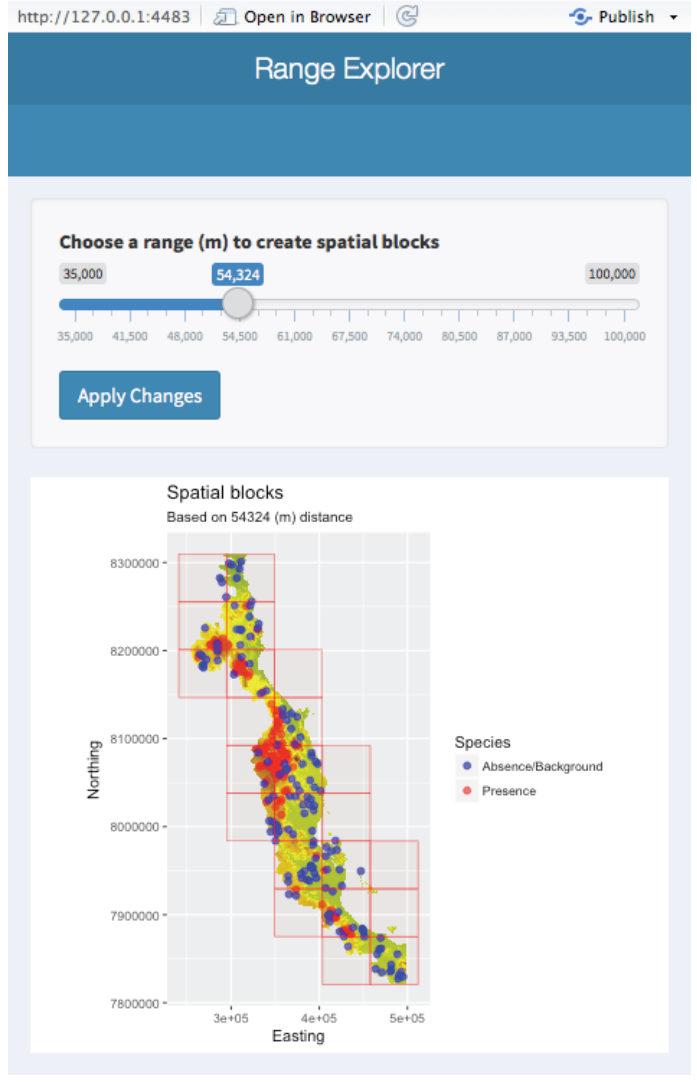
The graphical interface for choosing a suitable spatial block size

### Final remarks/Conclusion

The R package **blockCV** offers a suite of tools for creating data folds via blocking for evaluation of species distribution models, with three different blocking strategies, and tools that help users deal with typical nuances in species data, and for choosing block size and allocating blocks to folds. Which strategy is best to follow in a particular situation will depend on the purpose of the modelling and the region’s environmental structure (Roberts et al., 2017). For instance, both environmental blocks and horizontal spatial blocks could create distinct climatic groups in train and test datasets, which can be useful for assessing models aimed at predicting to new climatic conditions.

We recommend that users become familiar with the literature on block cross-validation, some of which we have cited in this article, so they can make appropriate choices. Different choices will have different implications on estimates of predictive performance. For instance, buffering is considered useful for enforcing spatial separation between train and test folds (Le Rest et al., 2014; Pohjankukka et al., 2017), but – depending on the relative sizes of the buffer and the region – it may produce training sets across repeats more similar than those produced by other blocking strategies. A disadvantage of similar training sets is that error estimates tend to have high variance (Hastie et al., 2009). Buffering also enforces as many training sets as presence or presence-absence points, so it can be quite computationally expensive. Other blocking strategies and choices of how to allocate blocks to folds will also have flow-on effects for estimates of performance, so it is important to think through what is most appropriate.

The package provides example data and a vignette with worked examples so users can explore the functions and learn how to use them in a species modelling workflow. The package will be actively maintained, and new features introduced as needs arise. We hope that this package enables modellers to more easily implement a range of evaluation approaches, so the modelling community learns more about the impacts of evaluation set-up on our understanding of predictive performance of SDMs.

## Availability

The package is available in GitHub (https://github.com/rvalavi/blockCV) and it will be available in CRAN and requires R version 3.4 or newer.

## Authors’ contributions

RV conceived the idea, wrote the code, drafted all documentation, and led the writing of the manuscript. All authors contributed to the design and testing of the package, and contributed critically to the manuscript drafts.

## Acknowledgments

RV is supported by an Australian Government Research Training Program Scholarship and a Rowden White Scholarship; GGA by an Australian Research Council (ARC) Discovery Early Career Researcher Award (DE160100904), and JJLM and JE by ARC Discovery Project 160101003. JE appreciates the support of the ARC’s Centre of Excellence for Environmental Decisions (CE11001000104). We also thank Babak Mirbagheri and Nick Golding for their helpful advice.

